# CagA delivery by the *Helicobacter pylori* Cag-Type IV Secretion System confers fitness benefits and costs during stomach infection

**DOI:** 10.64898/2026.08.07.743500

**Authors:** Jazmine A. Snow, Jacob P. Frick, Valerie P. O’Brien, Cynthia X. Guo, Scott D. Gray-Owen, Nina R. Salama

## Abstract

*Helicobacter pylori* strains encoding the *cag-*pathogenicity island (*cag-*PAI) and the effector toxin *cagA* are associated with worse disease outcomes. The *cag-*PAI encodes the Cag type IV secretion system (Cag-T4SS) which injects CagA and other bacterial products into gastric epithelial cells. Prior work revealed that host adaptive immunity promotes recombination in the *cag*-PAI gene *cagY* to attenuate Cag-T4SS activity during chronic infection, suggesting a fitness cost to assembling an active Cag-T4SS. To explore potential selective benefits and costs for the Cag-T4SS and CagA, we employed single strain and competitive infections at both acute and chronic timepoints in wildtype mice and transgenic mice that either attenuate innate immune responses or promote gastric pathology independent of *H. pylori* infection to examine the relative fitness of mutant *H. pylori* strains. Our results suggest that an active Cag-T4SS and CagA confer a fitness benefit during initial colonization through Cag-T4SS activity-dependent epithelial cell interactions that activate cancer-related signaling pathways. However, increasing gastric inflammation confers a fitness cost to CagA translocation, promoting Cag-T4SS shutoff. Targeted and whole genome sequencing revealed multiple mechanisms of Cag-T4SS attenuation, with recombination-mediated changes in *cagY* prevalent at early timepoints and mutations in a variety of Cag-T4SS structural genes accumulating as disease progresses. The need for Cag-T4SS activity and CagA translocation during initial gland colonization likely underlies the mutational pattern observed. Collectively this work reveals new insights into selective constraints on the *H. pylori* Cag-T4SS as well as resultant genetic adaptation processes that lead to retention of the *cag*-PAI and virulence.

## INTRODUCTION

*Helicobacter pylori* (*H. pylori*) infects roughly 44% of the global population, with ∼10% of infected individuals developing ulcers and 1-3% developing gastric cancer (1–5). Disease risk correlates in part with *H. pylori* genetic variation. The *cag*-pathogenicity island (*cag-*PAI) and *cagA,* which are present in 60-70% of *H. pylori* strains worldwide, have been associated with worse disease outcomes (6–14). The *cag*-PAI is a 40kb region that contains around 27 genes, with 17 of these being required for Cag-type IV secretion system (Cag-T4SS) activity along with the gene encoding the only known protein effector, CagA (15–19). The Cag-T4SS spans the inner and outer membrane of the bacterium and secretes CagA along with bacterial metabolites such as lipopolysaccharide (LPS) precursors, peptidoglycan fragments, and bacterial DNA into gastric epithelial cells (20–26). Once inside the gastric epithelial cells, CagA is phosphorylated by host kinases (SRC and ABL) and interacts with tyrosine phosphatase SHP2 (21, 27–34). CagA can manipulate numerous host cell pathways through phosphorylation-dependent and -independent mechanisms, promoting carcinogenesis (35–41). The other bacterial substrates secreted by the Cag-T4SS also impact host cells. LPS precursors (ADP-*glycero-*β-D-*manno-*heptose and heptose-1,7-bisphosphate) can be recognized by ALPK1 and lead to formation of TIFA-somes, resulting in the activation of NF-κB (23, 42–46). Peptidoglycan fragments activate NF-κB after recognition by NOD1 (24, 47, 48). CagA can also activate the NF-κB pathway, however the extent of activation varies based on the *H. pylori* strain and timepoint tested (49, 50). These distinct pathways of activating NF-κB in gastric epithelial cells all result in the production of IL-8 and other cytokines that recruit neutrophils and other immune cells to the site of infection (51–54).Experiments using gastric cancer cell lines have found that TIFA, NOD1, and then CagA sequentially activate NF-κB (55), however it is unclear whether a similar hierarchy occurs *in vivo*.

The Cag-T4SS can toggle between on and off states during infection (56, 57) primarily through genetic variation in *cagY*, encoding a member of the outer membrane core complex of the Cag-T4SS (56, 58–61). CagY has homology to VirB10 from other bacterial species at its C-terminus but is significantly larger than the canonical *virB10* from *Agrobacterium tumefaciens*, containing an intrinsically disordered 5’ repeat region and a large middle repeat region (MRR) comprised of two repeated motifs (A and B) (8, 62, 63). Recombination-mediated changes in the *cagY* MRR can increase or decrease the activity of the Cag-T4SS and has been suggested to serve as an immune rheostat triggered by host adaptive immune responses (64, 65).

While the Cag-T4SS and CagA promote disease, their roles in stomach colonization and persistence are less clear. All known *H. pylori* lineages except HpAfrica2 contain the *cag*-PAI at variable prevalence, with some lineages having near 100% *cag*-PAI positivity (66, 67). On the other hand, Cag-T4SS activity is often lost during human infection (65, 68) and in all animal models of stomach infection tested (56, 57, 60), suggesting selective pressures against sustained Cag-T4SS activity. In this study, we sought to clarify selective constraints on the Cag-T4SS during colonization, the role of its diverse substrates on measured selective constraints, and the impact of its activity across different stages of gastric disease. Our data suggest context-dependent selective pressures that depend on both bacterial metabolites and CagA.

## RESULTS

### Encoding the Cag-T4SS has context-dependent fitness benefits and costs during stomach infection

Prior observations of Cag-T4SS shutoff suggest a fitness cost that might only manifest after the onset of adaptive immunity (56, 65). To evaluate this, we performed single strain infection experiments for 6 weeks and 12 weeks, timepoints after the onset of the adaptive immune response in wildtype C57BL/6NJ mice following *H. pylori* infection. We investigated the impact of the Cag-T4SS on stomach colonization by infecting mice with PMSS1 Δ*cagE* (Δ*cagE*), a deletion mutant of *cagE* (an ATPase required for assembly of the Cag-T4SS (69, 70)) in *H. pylori* strain PMSS1 (our wildtype control strain, WT). PMSS1 is a *cag-*PAI+ strain known to encode an active Cag-T4SS and to robustly colonize the mouse stomach (71). In accordance with previous observations (72), Δ*cagE* showed a roughly one log increase in CFU per gram of stomach compared to WT after six weeks (median: 2.9 x 10^6^ Δ*cagE* vs. 1.6 x 10^5^ WT CFU/g stomach tissue) (**Figure 1A left**). At twelve weeks, Δ*cagE* continued to show higher colonization loads (median: 3.3 x 10^5^ Δ*cagE* vs. 9.1 x 10^4^ WT CFU/g stomach tissue) but to a lesser degree (**Figure 1A right**). To see if these changes in colonization were due to differences in growth rates, we cultured strains in liquid media and plated for CFU at timepoints between 0 and 28 hours. There were no differences in growth between WT and Δ*cagE* during liquid culture *in vitro* (**Supplemental Figure S1A**).

**Fig 1.**
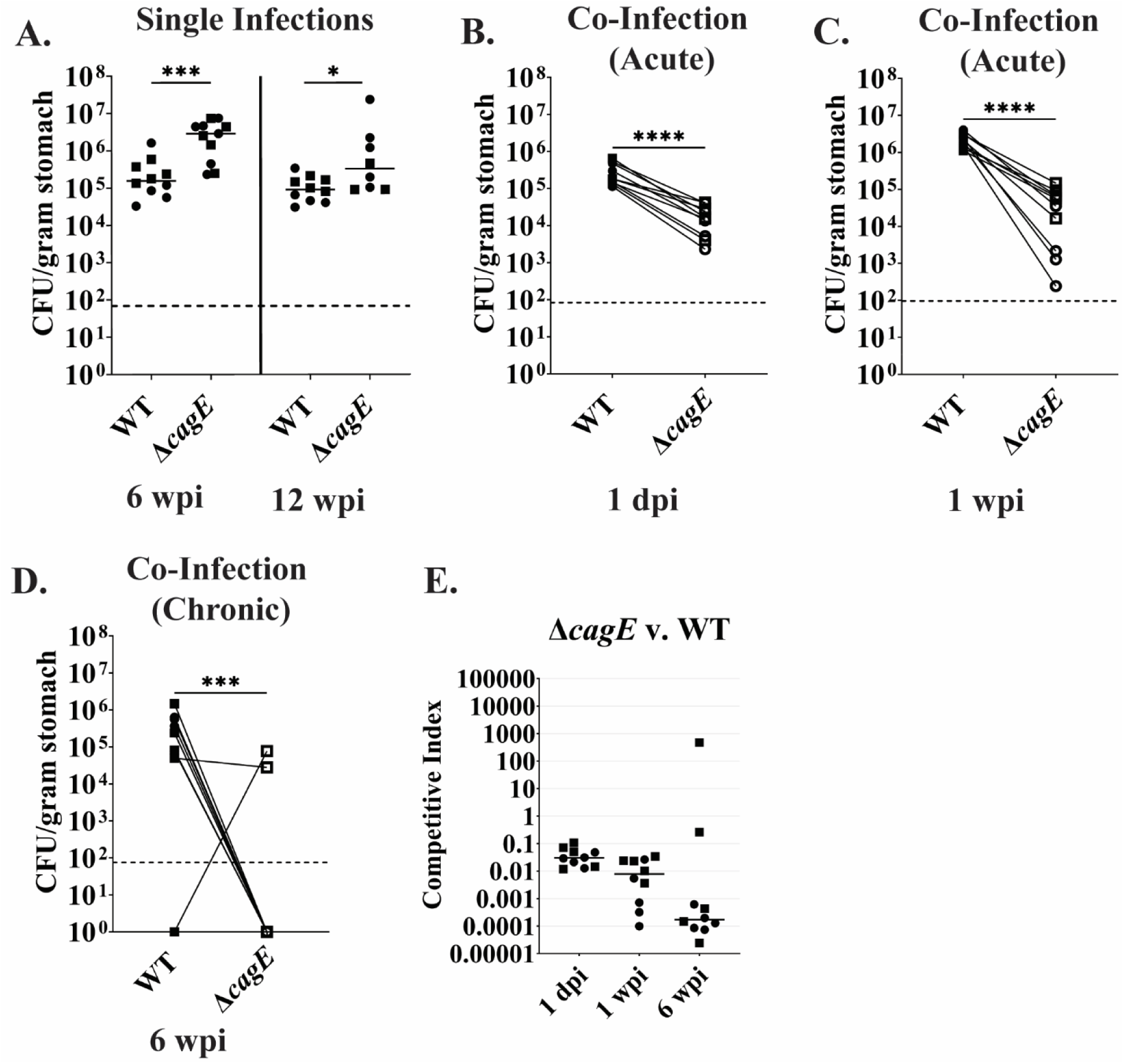
Encoding the Cag-T4SS has a fitness cost during single infections but has a fitness benefit during competitive infections. **A.** Colony forming units per gram of stomach tissue (CFU/gram stomach) for WT and Δ*cagE* after colonization in wildtype C57BL/6NJ mice for 6 weeks (**left**) or 12 weeks (**right**). Horizontal line depicts median CFU/g stomach for each strain. **B-D.** CFU/g stomach of WT and Δ*cagE* after co-infection in wildtype C57BL/6NJ mice for 1 day (**B**), 1 week (**C**), or 6 weeks (**D**). The lines connecting the points distinguish WT (solid symbols) and Δ*cagE* (open symbols) titers from the same mouse. Dotted line represents the limit of detection. **E.** Competitive index scores of B-D. Horizontal line depicts median competitive index score for each time point. dpi: days post infection. wpi: weeks post infection. N= 8-11 mice, 2 independent replicates differentiated by symbol shape. Nonparametric two-tailed Mann Whitney U test. *: p-value ≤ 0.05, ***: p-value ≤ 0.001, ****: p-value ≤ 0.0001.

For a more sensitive measurement of differential fitness, we inoculated mice with a 50:50 mixture of WT and Δ*cagE.* To examine initial colonization between the strains, we harvested stomachs for titers at 1 day and 1 week post infection using differential carriage of an antibiotic resistance cassette to distinguish among genotypes during plating. We also harvested stomachs at 6 weeks post infection to compare to differences observed during single infections at that time point. Unexpectedly, WT was able to significantly outcompete Δ*cagE* as early as 1 day post infection (p-value: <0.0001, nonparametric two-tailed Mann Whitney U test) (**Figure 1B, E**). At 1 week post infection, this competitive advantage increased with WT having roughly 1.6 logs higher median CFU/g of stomach (p-value: <0.0001, nonparametric two-tailed Mann Whitney U test) (**Figure 1C, E**). By 6 weeks post infection, 8/10 mice had cleared Δ*cagE* while retaining their WT infection (p-value: 0.0005, nonparametric two-tailed Mann Whitney U test) (**Figure 1D, E)**. We tested possible growth differences of 50:50 mixtures of WT and Δ*cagE* in liquid culture and plated for CFU at timepoints between 0 and 28 hours and found no differences in growth rates (**Supplemental Figure S1B**). To further test the possibility of growth differences, equal mixtures of WT and Δ*cagE* in liquid media were passaged daily for 3 days, plating for CFU each day. Again, we did not see any differences in growth between Δ*cagE* and WT *in vitro* (**Supplemental Figure S1C**). Thus, encoding a Cag-T4SS gave WT a slight fitness cost during single strain infections, but allowed WT to significantly outcompete Δ*cagE* during mixed infections. Furthermore, the selective advantage of the Cag-T4SS during competitive infection manifests before the onset of adaptive immunity.

### The Cag-T4SS effector CagA has differential effects on colonization before and after the onset of adaptive immunity

*H. pylori* induces an innate immune response early in colonization, eventually leading to the activation of the adaptive immune response and subsequent chronic inflammation within the host (73, 74). Prior work suggested that induction of innate immune responses is largely CagA-independent and is instead driven by the translocation of bacterial metabolites by the Cag-T4SS. However, CagA can contribute to NF-κB activation in select strains of *H. pylori* (24, 46, 49). CagA additionally induces several other signaling pathways in host cells, including MAP kinase (MAPK) signaling and β-catenin activation (41, 75–77). *H. pylori* strain PMSS1 may contain multiple copies of *cagA* (typically between 0-4) with the copy number expanding and contracting during infection (78, 79). To clarify which substrates, and thus possible host cell pathways, underlie selective pressures on Cag-T4SS activity, we tested a mutant version of PMSS1 that lacks all copies of *cagA* (Δ*cagA*) while still encoding a functional Cag-T4SS that can secrete the other Cag-T4SS substrates, in single strain and competitive infections. At both six and twelve-weeks post infection, mice infected with Δ*cagA* showed slightly higher levels of colonization than WT in healthy stomach tissue (6 weeks post infection p-value: 0.13, 12 weeks post infection p-value: 0.11, nonparametric two-tailed Mann Whitney U test) (**Figure 2A**). To see if these changes in colonization were due to differences in growth rates, we cultured strains in liquid media and plated for CFU at timepoints between 0 and 28 hours and found no differences (**Supplemental Figure S2A**). Unlike the WT/Δ*cagE* competitive infections, Δ*cagA* slightly outcompeted WT at 1 day post infection (p-value: 0.01, nonparametric two-tailed Mann Whitney U test) (**Figure 2B, E**). However, the opposite was seen at 1 week post infection, where WT colonized at significantly higher loads than Δ*cagA* (p-value: <0.0001, nonparametric two-tailed Mann Whitney U test) (**Figure 2C, E**). After six weeks, the colonization was mixed, with most of the mice having either equal levels of WT and Δ*cagA* (4/10) or having WT outcompete Δ*cagA* (4/10) (**Figure 2D, E**). In two of the mice Δ*cagA* had outcompeted WT (2/10). Like Δ*cagE,* we found no differences in growth compared to WT during co-culture of Δ*cagA* and WT in liquid media for up to 3 days (**Supplemental Figure S2B, C**). These results suggest that while the presence of CagA plays a role in the competitive fitness advantage seen for WT, other Cag-T4SS substrates likely contribute before the onset of adaptive immunity (six weeks), after which the competitive advantage of possessing CagA appears reduced and may even confer a fitness cost.

**Fig 2.**
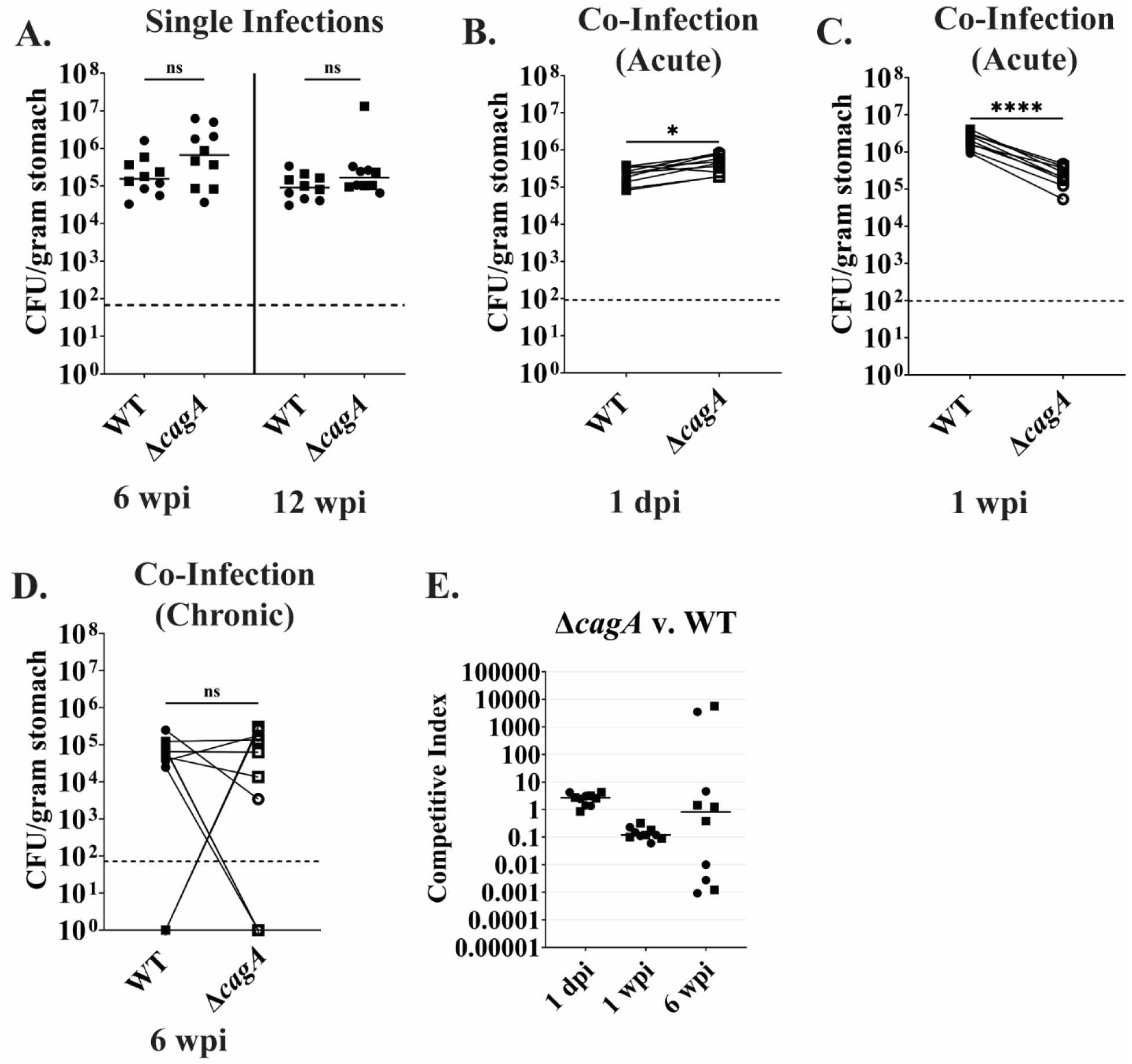
CagA confers mixed fitness phenotypes, playing a larger role before the onset of the adaptive immune response. **A.** Colony forming units per gram of stomach tissue (CFU/gram stomach) for WT and Δ*cagA* after colonization in wildtype C57BL/6NJ mice for 6 weeks (**left**) or 12 weeks (**right**). Horizontal line depicts median CFU/g stomach for each strain. **B-D.** CFU/g stomach of WT and Δ*cagA* after co-infection in wildtype C57BL/6NJ mice for 1 day (**B**), 1 week (**C**), or 6 weeks (**D**). The lines connecting the points distinguish WT (solid symbols) and Δ*cagA* (open symbols) titers from the same mouse. Dotted line represents the limit of detection. **E.** Competitive index scores of B-D. Horizontal line depicts median competitive index score for each timepoint. dpi: days post infection. wpi: weeks post infection. N= 10 mice, 2 independent replicates differentiated by symbol shape. Nonparametric two-tailed Mann Whitney U test. ns: nonsignificant, *: p-value ≤ 0.05, ****: p-value ≤ 0.0001.

### Innate immune sensors TIFA and NOD1 do not impact colonization patterns seen in wildtype C57BL/6NJ mice

Since the colonization advantage seen at the earliest timepoints appeared to be independent of CagA presence, we examined whether the presence of TIFA or NOD1, known innate immune sensors of bacterial metabolites delivered by the Cag-T4SS (23, 24, 42, 46), modulate colonization by *H. pylori*. Mice lacking either *Nod1, Tifa,* or both (*Tifa/Nod1* knock-out mice) were infected with a 50:50 mixture of WT and Δ*cagE.* WT outcompeted Δ*cagE* at 1 day post infection regardless of *Tifa* or *Nod1* status (**Figure 3A, B**). The competitive advantage was even greater by 1 week of infection (**Figure 3C, D**). By 6 weeks post infection, most mice (16/20) had cleared Δ*cagE* while retaining WT, irrespective of their *Tifa* or *Nod1* genotypes (**Figure 3E, F**). Moreover, overall bacterial loads for each strain were comparable in all mice, regardless of *Tifa* or *Nod1* genotype. To further clarify whether activation of TIFA impacts colonization in general and in the absence of CagA, competition experiments between WT and Δ*cagA* were repeated in mice containing two (*Tifa* +/+), one (*Tifa* +/-), or zero (*Tifa* -/-) *Tifa* alleles. At 1 day post infection, we saw no differences in WT or Δ*cagA* titers between *Tifa* +/+, *Tifa* +/-, or *Tifa* -/- mice, and regardless of *Tifa* presence, there were relatively equal levels of WT and Δ*cagA* (**Supplemental Figure S3A, B**). WT outcompeted Δ*cagA* by about 1.5 to 2 logs at 1 week post infection regardless *Tifa* presence (*Tifa* +/+ (p-value: 0.0002), *Tifa* +/- (p-value: 0.0022), *Tifa* -/- (p-value: 0.0006), nonparametric two-tailed Mann Whitney U test) (**Figure 3G, H**). Results were mixed at 6 weeks post infection with 6/9 *Tifa* +/+, 6/12 *Tifa* +/-, and 8/11 *Tifa* -/- mice having cleared Δ*cagA* while retaining their WT infection. The remaining mice either had roughly equal levels of WT and Δ*cagA* or more rarely, only Δ*cagA* (**Supplemental Figure S3C, D**). Taken together, colonization levels and the competitive infection results for WT and Δ*cagA* mirrored those seen in wildtype C57BL/6NJ mice in the prior experiments. These results suggest that Cag-T4SS-dependent bacterial cell envelope metabolite delivery does not play a major role in the selective advantage of encoding the Cag-T4SS during initial colonization, nor does it influence the frequency at which *cagA* mutants appear favored later in infection.

**Fig 3.**
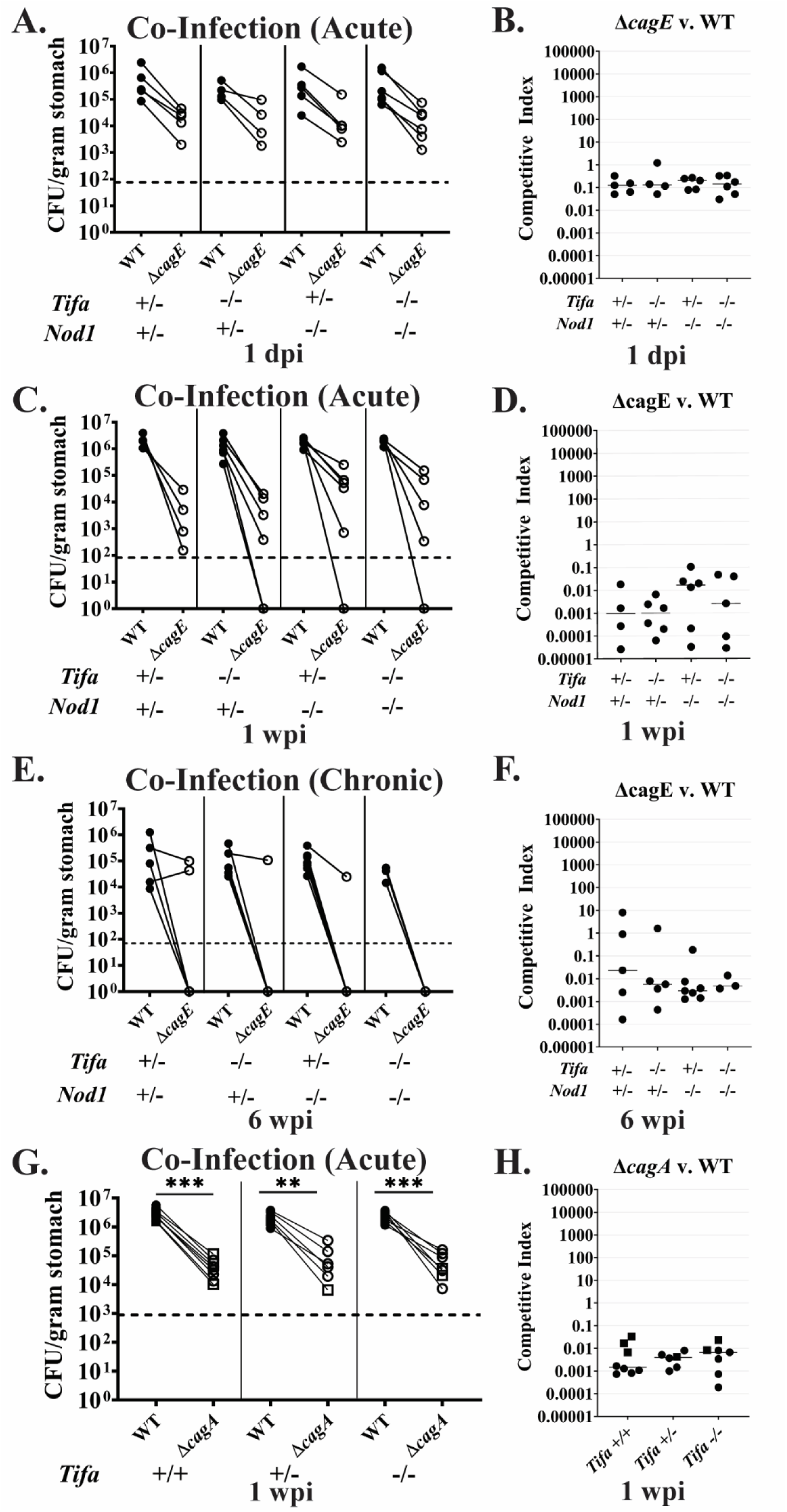
Host *Tifa* and *Nod1* status does not impact overall colonization or competition outcomes during stomach infection. **A.** Colony forming units per gram of stomach tissue (CFU/gram stomach) for WT and Δ*cagE* after 1 day of colonization in mice with the indicated *Tifa* and *Nod1* genotypes. **B.** Competitive index scores of A. **C.** CFU/gram stomach for WT and Δ*cagE* after 1 week of colonization in mice with the indicated *Tifa* and *Nod1* genotypes. **D.** Competitive index scores of C. **E.** CFU/gram stomach for WT and Δ*cagE* after 6 weeks of colonization in mice with the indicated *Tifa* and *Nod1* genotypes. The lines connecting the points distinguish WT (solid circles) and Δ*cagE* (open circles) titers from the same mouse. **F.** Competitive index scores of E. **G.** CFU/gram stomach for WT and Δ*cagA* after 1 week of colonization in mice with two (*Tifa +/+*), one (*Tifa +/-*), or no (*Tifa -/-*) copies of *Tifa.* The lines connecting the points distinguish WT (solid symbols) and Δ*cagA* (open symbols) titers from the same mouse. Nonparametric two-tailed Mann Whitney U test. **: p-value ≤ 0.01, ***: p-value ≤ 0.001. Dotted line represents the limit of detection. **H.** Competitive index scores of G. N= 3-8 mice, 1-2 independent replicates differentiated by symbol shape.

### CagA promotes Cag-T4SS shutoff during stomach infection

While our WT/Δ*cagE* competitive infections showed a beneficial role for encoding the Cag-T4SS during colonization (**Figure 1**), our WT/Δ*cagA* competitive infections varied by timepoint (**Figure 2**). These results led us to wonder whether sustained CagA translocation may contribute to the selective pressures that lead to Cag-T4SS shutoff during chronic infection. To assess this, we recovered WT or Δ*cagA* output strains after colonization and indirectly determined their relative Cag-T4SS activity. To do so, we co-cultured *H. pylori* strains isolated from mice with AGS cells (a human gastric cancer cell line) for 24 hours and measured the production of IL-8, which is dependent on a functional Cag-T4SS. We tested approximately 7-8 output strains from 2-3 mice per experiment and normalized IL-8 induction against their relative input controls (WT input pre-mouse infection for WT output strains and Δ*cagA* input pre-mouse infection for Δ*cagA* output strains). It is important to note that Δ*cagA* induces less IL-8 compared to WT; however, it is still able to induce IL-8 due to the presence of the non-protein Cag-T4SS substrates. Representative unnormalized data are shown in **Supplemental Figure S4**.

We first examined output strains after 6 weeks of infection, which is after the induction of adaptive immunity. WT output strains induced significantly less IL-8 in AGS cells relative to their input strain than the Δ*cagA* output strains (median 58% vs. 82%, p-value: <0.0001, nonparametric two-tailed Mann Whitney U test) (**Figure 4A left**), suggesting that the presence of CagA was important in Cag-T4SS shutoff at this timepoint.

**Fig 4.**
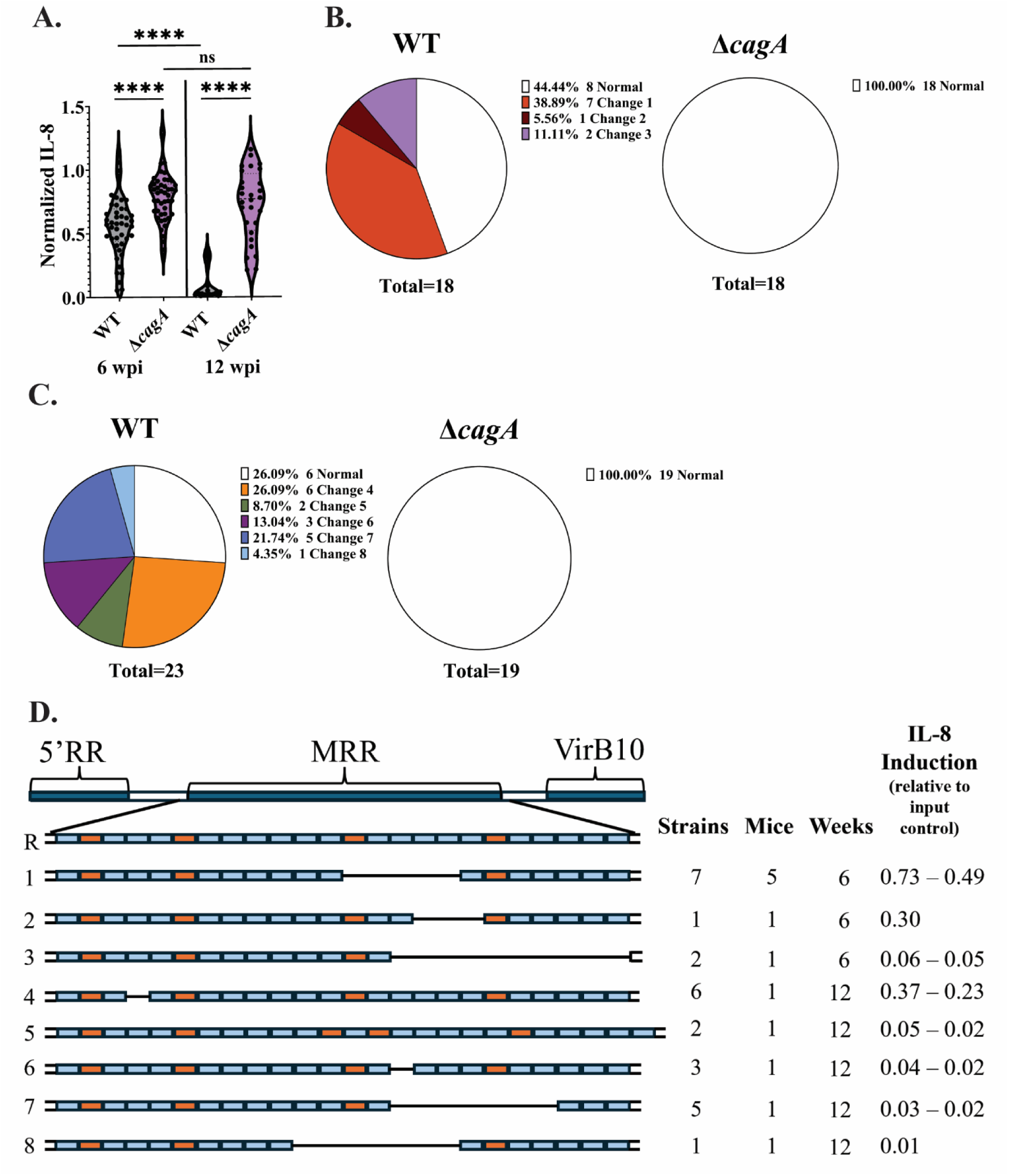
CagA drives Cag-T4SS shutoff during colonization in wildtype C57BL/6NJ mice, often by recombination in CagY. **A.** Levels of IL-8 induction in AGS cells after co-culture with WT or Δ*cagA* output strains after 6 weeks (left) or 12 weeks (right) of infection in wildtype C57BL/6NJ mice, relative to their respective input controls. Each point represents an individual output strain with 28-47 output strains taken from 4-6 mice from 2 independent experiments per bacterial strain. Nonparametric two-tailed Mann Whitney U test. ns: nonsignificant, ****: p-value ≤ 0.0001. **B.** Percentage of output strains with altered *cagY* MRR for WT and Δ*cagA* output strains after 6 weeks of infection in wildtype C57BL/6NJ mice. **C.** Percentage of output strains with normal or changed *cagY* MRR for WT and Δ*cagA* output strains after 12 weeks of infection in wildtype C57BL/6NJ mice. **D.** Schematic representing the different changes seen in the *cagY* MRR in WT output strains after 6 or 12 weeks of infection in C57BL/6NJ mice. 5’RR= 5’ repeat region, MRR= middle repeat region, VirB10= *virB10* homology region. Blue= A motifs, Orange= B motifs. R= reference *cagY* MRR. Numbers 1-8 on the left each represent a different change. Strains= number of different output strains that had the change. Mice= the number of different mice that had output strains with that change. Weeks= weeks postinfection from which output strains were recovered. IL-8 Induction= the relative IL-8 induction of output strains had compared to input control (value or range).

Prior work identified sequence variation in *cagY*, encoding a member of the Cag-T4SS outer membrane core complex, as an important regulator of Cag-T4SS activity during infection (56, 64). Specifically, recombination within the middle repeat region (MRR) of *cagY* can result in differences in the number or pattern of A and B motifs comprising the MRR, impacting Cag-T4SS activity (56, 62, 64). To assess if CagY variation accounted for lower IL-8 induction in these output strains, we sequenced the MRR of *cagY* in WT output strains that exhibited a range of IL-8 induction. Overall, 10 of 18 WT output strains sequenced had recombination resulting in in-frame deletions within their *cagY* MRR compared to 0 of the 18 Δ*cagA* output strains tested (**Figure 4B**).

We tested additional output strains obtained after 12 weeks of colonization to examine whether Δ*cagA* output strains would eventually lose Cag-T4SS activity. Consistent with previous findings (56, 65), WT output strains from 12 weeks induced less IL-8 compared to the WT output strains from 6 weeks of infecting wildtype C57BL/6NJ mice (median 0.02% vs. 58%, p-value: <0.0001, nonparametric two-tailed Mann Whitney U test) (**Figure 4A**). However, the difference in IL-8 induction between Δ*cagA* output strains after 6 or 12 weeks of colonization was nonsignificant (median 82% vs. 77%, p-value: 0.565602, nonparametric two-tailed Mann Whitney U test) (**Figure 4A**). While most of the WT output strains (21/28) had complete shutoff (having induced less than 10% of the IL-8 seen in the WT input control), with a median IL-8 induction at 0.02% relative to input controls, the Δ*cagA* output strains retained Cag-T4SS activity, with a median IL-8 induction of 77% compared to the Δ*cagA* input control (**Figure 4A right**). Again, WT had significantly more Cag-T4SS shutoff (p-value: <0.0001, nonparametric two-tailed Mann Whitney U test), suggesting that the presence of CagA is the primary Cag-T4SS substrate driving shutoff during chronic infection. Changes within *cagY* were also seen in WT output strains at 12 weeks post infection, with 17 of the 23 WT output strains having alterations in their *cagY* MRR. Again, 0 of the 19 Δ*cagA* output strains sequenced had changes in their *cagY* MRR regardless of their ability to induce IL-8 in AGS cells (**Figure 4C**). Therefore, even at this longer time point, Δ*cagA* output strains continue to induce IL-8 and regulation of their Cag-T4SS activity is largely *cagY* recombination-independent.

To determine if there were any distinct patterns in *cagY* MRR alterations that impact Cag-T4SS activity in these output strains, we mapped the different alterations to the *cagY* MRR that were present within our 6- and 12-week WT output strains that had changes in *cagY* (**Figure 4D)**. In the 6 weeks post infection output strains, Change #1, consisting of a 549 bp in-frame deletion resulting in the loss of four A motifs and one B motif, was found the most often, having occurred in seven output strains among five separate mice, and conferred a range of IL-8 induction from 78% to 49% of WT input control levels. Change #2, with a 345 bp in-frame deletion leading to the loss of three A motifs, only occurred in one output strain leading to 30% IL-8 induction compared to WT input control levels. Change #3 occurred in two output strains recovered from the same mouse that had complete Cag-T4SS shut off, inducing less than 10% of the IL-8 induced by the input control. This change was a large 1,221 bp in-frame deletion resulting in the loss of nine A motifs and one B motif (**Figure 4D**).

The output strains recovered after 12 weeks of infection in wildtype C57BL/6NJ mice had unique alterations in their *cagY* MRR compared to the 6-week output strains (**Figure 4D**). Change #4 occurred in output strains ranging from 37% to 23% relative IL-8 induction compared to WT input control levels and was a small 117 bp in-frame deletion resulting in the loss of one A motif. The rest of the output strains sequenced had completely shut off the Cag-T4SS, with less than 10% IL-8 induction compared to WT input control. Rather than an in-frame deletion, Change #5 comprised an extra B motif. Change #6 was a small 114 bp in-frame deletion resulting in the loss of one A motif. Change #7 was a large 780 bp in-frame deletion leading to the loss of six A motifs and one B motif. Change #8 was a large 777 bp in-frame deletion with a loss of six A motifs and one B motif. Output strains with Changes 5, 6, and 8 all came from the same mouse, while the output strains with other *cagY* MMR changes each came from a unique mouse. Despite finding a range of alterations in the *cagY* MRR for output strains taken 6 or 12 weeks after colonization in healthy C57BL/6NJ mice, we could not find any patterns between the MRR changes and the degree of Cag-T4SS shut off.

Not all WT output strains with lower Cag-T4SS activity and none of the Δ*cagA* output strains sequenced had changes in the *cagY* MRR, despite detectable differences in the level of IL-8 induction compared to their input control. To assess other changes in the *cag-*PAI and beyond, we performed Oxford Nanopore long-read whole genome sequencing (WGS) of select WT and Δ*cagA* output strains after 6 weeks of colonization representing a range of IL-8 induction and *cagY* MRR changes from different mice (results summarized in **Table 1)**. One WT output strain we sequenced, Mouse A WT output (MsA WT) had similar levels of IL-8 induction as the input control (Relative IL-8 induction: 114%) and had no detectable *cag-*PAI changes compared to the WT input control. We then sequenced MsB WT, which had moderate attenuation of Cag-T4SS activity (Relative IL-8 induction: 55%) and Change #1 in its *cagY* MRR. WGS revealed that the recombination in *cagY* was the only alteration in the *cag-*PAI for this output strain We also sequenced two WT output strains that had decreased IL-8 induction (Relative IL-8 induction: 24% and 12%) but no changes in the *cagY* MRR. Both strains had fewer copies of *cagA* compared to the WT input control (which typically has four copies (78)). This is in line with previous work that has shown that PMSS1 can expand and contract its *cagA* copy number both *in vivo* and *in vitro,* impacting its ability to induce IL-8 in AGS cells (78). MsC WT only had two *cagA* copies. MsD WT had three copies of *cagA* and had a frameshift mutation in *cagW*, encoding a VirB6 homologue required for Cag-T4SS activity.

**Table 1:** Alterations to *cag-*PAI genes found in output strains after six weeks of infection in wildtype C57BL/6NJ mice.

| Output strain | Locus | Vir gene homology | Polymorphism | Type | IL-8 Induction<sup>a</sup> | CagY MRR<sup>b</sup> |
| --- | --- | --- | --- | --- | --- | --- |
| WT output strains |  |  |  |  |  |  |
| MsA WT |  |  | Same as input |  | 1.14 | NA |
| MsB WT | cagY | VirB10 | MRR variation | Repeat recombination | 0.55 | Change 1 |
| MsC WT | cagA |  | ΔcagA_1_cagA_2 | Copy number variation | 0.24 | NC |
| MsD WT | cagW | VirB6 | +1bp coding (10/759) | Frameshift | 0.12 | NC |
|  | cagA |  | ΔcagA_1 | Copy number variation |  |  |
| ΔcagA output strains |  |  |  |  |  |  |
| MsE ΔcagA | cag5 | VirD4 | +1bp coding (271/2247) | Frameshift | 1.32 | NC |
| MsF ΔcagA |  |  | Only has ΔcagA_1-4, same as input |  | 1.10 | NC |
| MsG ΔcagA |  |  | Only has ΔcagA_1-4, same as input |  | 0.88 | NC |
| MsH ΔcagA |  |  | Only has ΔcagA_1-4, same as input |  | 0.76 | NC |
| MsI ΔcagA | cagT | VirB7 | Δ1bp coding (780/843) | Frameshift | 0.48 | NC |

<sup>a</sup>IL-8 induction relative to respective input controls.
<sup>b</sup>Results of *cagY* MRR sequencing. NA: not applicable, output strain did not have *cagY* MRR sequenced. NC: no change, *cagY* MRR was the same as input control. Change #: the change seen from *cagY* MRR sequencing (**Figure 4D**).

For the Δ*cagA* output strains, we sequenced two strains (MsE Δ*cagA* and MsF Δ*cagA*) that induced IL-8 at similar levels to input control (Relative IL-8 induction: 132% and 110%, respectively).) MsE Δ*cagA* had a frameshift mutation in *cag5,* encoding a VirD4 homologue required for CagA translocation. MsF Δ*cagA* had no detectable changes in the *cag-*PAI when compared to the Δ*cagA* input control. We also sequenced three Δ*cagA* output strains that had decreased IL-8 induction compared to the input control. Two output strains (MsG Δ*cagA* and MsH Δ*cagA*) had slightly decreased IL-8 induction (Relative IL-8 induction: 88% and 76%, respectively) but did not have any differences in their *cag-*PAI compared to input control. MsI Δ*cagA* induced moderate levels of IL-8 compared to its input control (Relative IL-8 induction: 48%) and had a frameshift mutation near the end of *cagT* encoding a VirB7 homologue required for Cag-T4SS activity. Collectively, we found that *cagY* MRR variation does occur in the context of WT infection along with other less common *cag-*PAI changes; however, in Δ*cagA,* shut off occurs mainly through CagY-independent mechanisms.

We also performed WGS on WT and Δ*cagA* output strains obtained after 12 weeks of infection (results summarized in **Table 2)**. In all cases, WGS confirmed the *cagY* MRR changes previously identified by amplicon sequencing. In addition, MsJ WT (Relative IL-8 induction: 37%) and MsL WT (Relative IL-8 induction: 1%) which had *cagY* MRR Change #4 and Change #8 respectively, both carried only one copy of *cagA.* MsK WT (Relative IL-8 induction: 2%), which had Change #7 in the *cagY* MRR, contained five copies of *cagA.* We also sequenced one output strain, MsM WT, that did not have differences in the *cagY* MRR but had complete Cag-T4SS shutoff (Relative IL-8 induction: 1%). WGS revealed two copies of *cagA* and a frameshift mutation in *cagD*, encoding CagD which has been found to have a partial impact on Cag-T4SS activity and CagA translocation (17, 20, 26, 80).

**Table 2:** Alterations to *cag-*PAI genes found in output strains after 12 weeks of infection in wildtype C56BL/6J mice.

| Output strain | Locus | Vir gene homology | Polymorphism | Type | IL-8 Induction<sup>a</sup> | CagY MRR<sup>b</sup> |
| --- | --- | --- | --- | --- | --- | --- |
| WT output strains |  |  |  |  |  |  |
| MsJ WT | *cagY* | VirB10 | MRR variation | Repeat recombination | 0.37 | Change 4 |
| | *cagA* | | Δ*cagA*_1_*cagA*_2_*cagA*_3 | copy number variation | | |
| MsK WT | *cagY* | VirB10 | MRR variation | Repeat recombination | 0.02 | Change 7 |
|  | *cagA* |  | +*cagA*_5 | copy number variation |  |  |
| MsL WT | *cagY* | VirB10 | MRR variation | Repeat recombination | 0.01 | Change 8 |
| | *cagA* | | Δ*cagA*_1_*cagA*_2_*cagA*_3 | copy number variation | | |
| MsM WT | *cagD* |  | +1bp coding 72/660 | Frameshift | 0.01 | NC |
| | *cagA* | | Δ*cagA*_1_*cagA*_2 | copy number variation | | |
| Δ*cagA* output strains | | | | | | |
| MsN Δ*cagA* | | | Δ*cagA*_1-4, same as input | | 1.16 | NC |
| MsO Δ*cagA* | | | Δ*cagA*_1-4, same as input | | 0.96 | NC |
| MsP Δ*cagA* | | | Δ*cagA*_1-4, same as input | | 0.86 | NC |
| MsQ Δ*cagA* | | | Δ*cagA*_1-4, same as input | | 0.78 | NC |

<sup>a</sup>IL-8 induction relative to respective input controls.
<sup>b</sup>Results of *cagY* MRR sequencing. NC: no change, *cagY* MRR was the same as input control.
Change #: the change seen from *cagY* MRR sequencing (**Figure 4D**).

We sequenced four Δ*cagA* output strains (MsN Δ*cagA*, MsO Δ*cagA*, MsP Δ*cagA* and MsQ Δ*cagA*) that had similar or slightly lower levels of IL-8 induction to input control (Relative IL-8 induction: 116-78%). WGS did not reveal any changes in the *cag-*PAI for any of these four strains. Taken together, alterations in both the *cagY* MRR and in other *cag-*PAI genes increased with longer infection time in WT output strains while Δ*cagA* output strains continue to have less Cag-T4SS shut off correlating with less changes in the *cag-*PAI.

### Encoding the Cag-T4SS confers context-dependent fitness benefits and costs during preneoplasia

Previous work has shown that specific virulence factors may be more important for colonization during preneoplasia of the stomach (81, 82), when gland organization and cell composition is markedly different than in healthy glands (83, 84). Therefore, we investigated whether encoding the Cag-T4SS impacts colonization in metaplastic stomach tissue. To do this, we utilized the *Mist1-Kras* (*Mist1-CreERT2^Tg/+^; LSL-K-Ras*(*G12D*)*^Tg/+^*) mouse model of gastric preneoplasia (85, 86). The *Mist1-Kras* mouse model develops metaplasia after expression of Kras^G12D^ is induced in *Mist1-*expressing cells (specifically, chief cells and a subset of isthmus progenitor cells in the stomach). To test the impact of the Cag-T4SS and its substrates in this model, we induced Kras^G12D^ using tamoxifen (TMX). By 6 weeks post Kras^G12D^ induction, these mice develop metaplasia with intestinal characteristics in the stomach, referred to herein as gastric intestinal metaplasia (GIM). At this point, *Mist1-Kras* mice with GIM were infected with WT or Δ*cagE* for an additional 6 weeks. In accordance with prior observations (87), Δ*cagE* had significantly elevated titers (median: 1.3 x 10^5^ CFU/g of stomach) compared to WT (median: 1.2 x 10^4^ CFU/g of stomach, p-value: 0.03, nonparametric two-tailed Mann Whitney U test), despite the broad range of colonization for each strain (**Figure 5A**). This suggests that encoding the Cag-T4SS when colonizing metaplastic stomach tissue still confers a fitness cost. To see if WT still outcompetes Δ*cagE* when both are colonizing the same stomach, as occurred in wildtype C57BL/6NJ mice, competitive infections were performed in *Mist1-Kras* mice with GIM. At 1 day post infection, WT outcompeted Δ*cagE* (p-value: 0.04, nonparametric two-tailed Mann Whitney U test) (**Figure 5B, E**) and similar to our observations during infection of wildtype C56BL/6NJ mice, the colonization advantage of WT over Δ*cagE* increased with the length of infection. WT outcompeted Δ*cagE* by 2.2 logs by 1 week post infection (p-value: <0.0001, nonparametric two-tailed Mann Whitney U test) (**Figure 5C, E**). By 6 weeks post infection, in all but one mouse, Δ*cagE* fell below the limit of detection while WT persisted. In the one mouse with Δ*cagE* remaining, WT had a 2-log increase in CFU/g stomach (p-value: <0.0001, nonparametric two-tailed Mann Whitney U test) (**Figure 5D, E**). Therefore, encoding the Cag-T4SS promotes colonization during competitive infection in *Mist1-Kras* mice with GIM, similar to when colonizing healthy stomach tissue of wildtype C57BL/6NJ mice.

**Fig 5.**
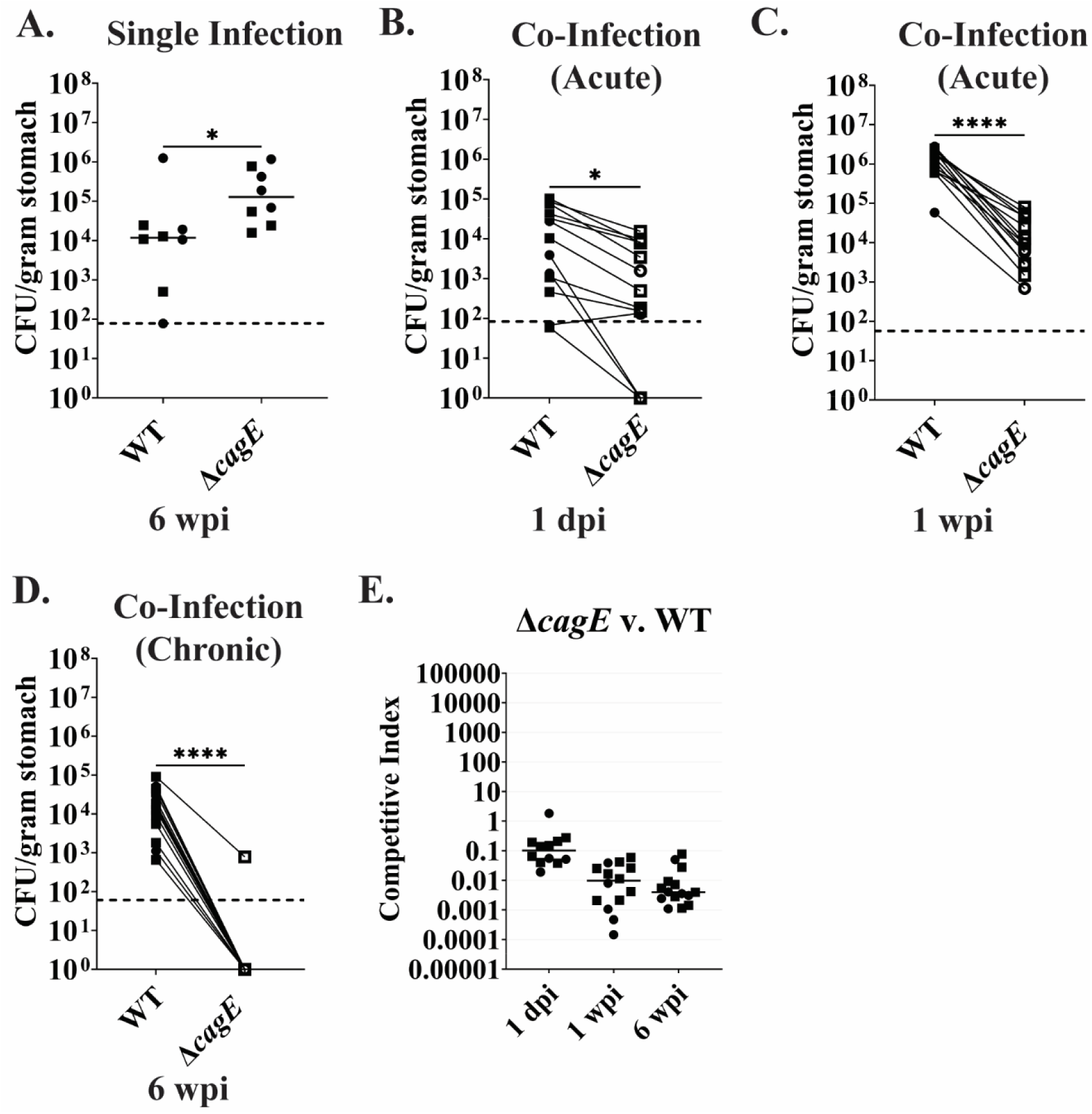
Encoding the Cag-T4SS continues to have a fitness cost during single infections and a colonization benefit during competitive infections in *Mist1-Kras* mice with GIM. *Mist1-Kras* mice received subcutaneous tamoxifen (TMX) injections on days 1-3 to induce Kras^G12D^ expression in *Mist1+* cells. 6 weeks after TMX administration, at which time gastric intestinal metaplasia (GIM) had developed, mice were infected with *H. pylori* via oral gavage. **A.** Colony forming units per gram of stomach tissue (CFU/gram stomach) for WT and Δ*cagE* after six weeks of infection in *Mist1-Kras* mice with GIM. Horizontal line depicts median CFU/g stomach for each strain. **B-D.** CFU/g stomach of WT and Δ*cagE* after co-infection in *Mist1-Kras* mice with GIM for 1 day (**B**), 1 week (**C**), or 6 weeks (**D**). The lines connecting the points distinguish WT (solid symbols) and Δ*cagE* (open symbols) titers from the same mouse. Dotted line represents the limit of detection. **E.** Competitive index scores of B-D. Horizontal line depicts median competitive index score for each timepoint. dpi: days post infection. wpi: weeks post infection. N= 8-15 mice, 2 independent replicates differentiated by symbol shape. Nonparametric two-tailed Mann Whitney U test. *: p-value ≤ 0.05, ****: p-value ≤ 0.0001.

### CagA effect on colonization is abrogated in GIM stomach tissue

We also tested the impact of CagA on colonization in the metaplastic stomach. After 6 weeks of colonization in *Mist1-Kras* mice with GIM, Δ*cagA* (median: 1.6 x 10^5^ CFU/g of stomach) tended to have higher loads compared to WT (median: 1.2 x 10^4^ CFU/g of stomach); however, the variability in titers made the trend nonsignificant (p-value: 0.06, nonparametric two-tailed Mann Whitney U test) (**Figure 6A**). We also repeated the competitive infections for WT and Δ*cagA* at 1 day, 1 week, and 6 weeks of colonization in *Mist1-Kras* mice with GIM. At one day post infection, WT and Δ*cagA* showed similar colonization levels (p-value: 0.13, nonparametric two-tailed Mann Whitney U test) (**Figure 6B, E**). Notably, 4/13 *Mist1-Kras* mice with GIM had cleared their WT infection while retaining Δ*cagA* at 1 day post infection. The levels of WT and Δ*cagA* were equal at 1 week post infection (p-value: 0.38, nonparametric two-tailed Mann Whitney U test) (**Figure 6C, E**) through 6 weeks post infection (p-value: 0.25, nonparametric two-tailed Mann Whitney U test) (**Figure 6D, E**). This differed from our observations in wildtype C57BL/6NJ mice where WT outcompeted Δ*cagA* beginning at 1 week (**Figure 2C, E**), Collectively, these results indicate that possessing CagA does not influence WT’s competitive advantage over Δ*cagE* when colonizing the metaplastic stomach, even at early time points.

**Fig 6.**
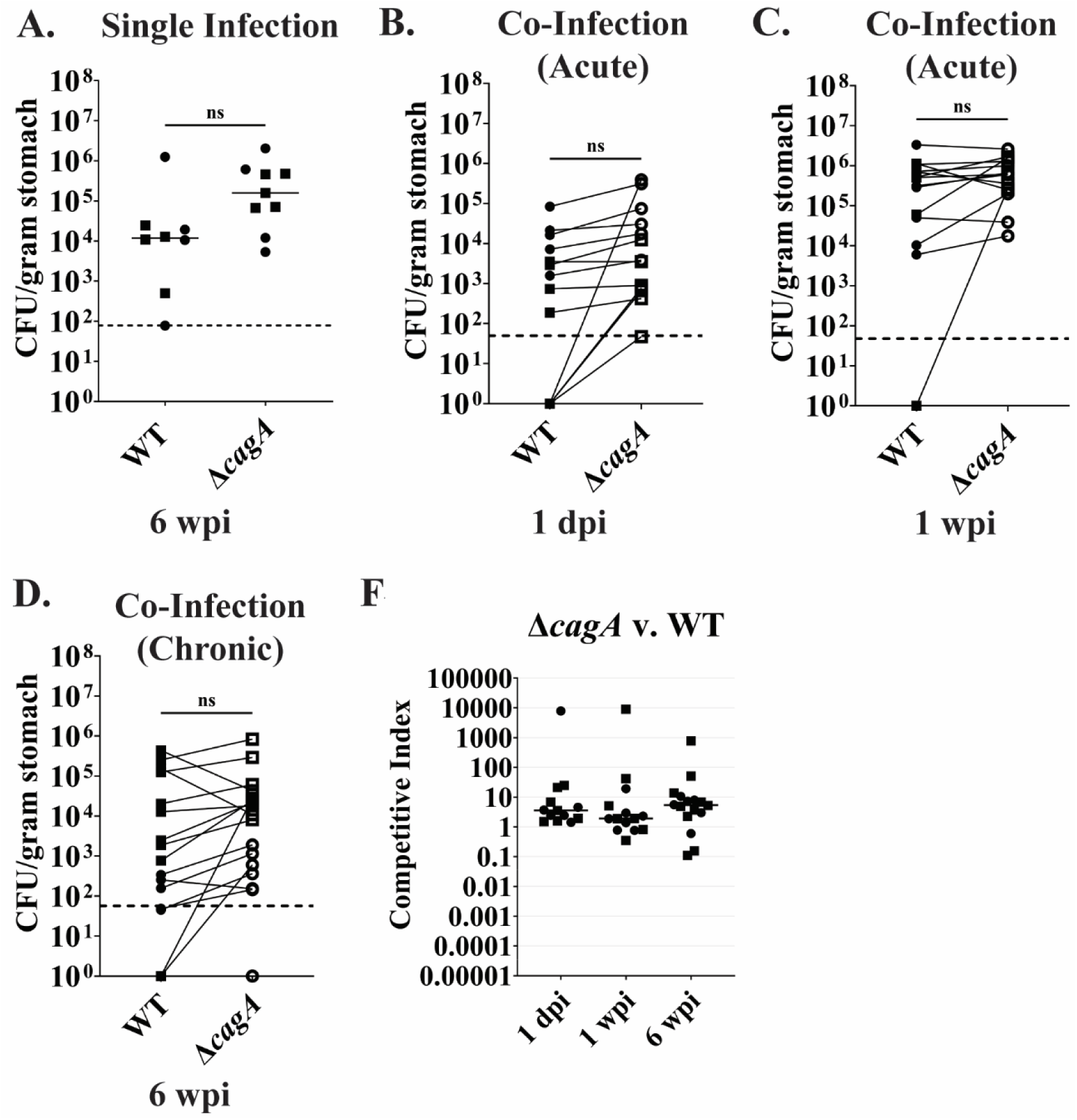
CagA does not impact colonization in *Mist1-Kras* mice with GIM. *Mist1-Kras* mice received subcutaneous tamoxifen (TMX) injections on days 1-3 to induce Kras^G12D^ expression in *Mist1+* cells. 6 weeks after TMX administration, at which time gastric intestinal metaplasia (GIM) had developed, mice were infected with *H. pylori* via oral gavage. **A.** Colony forming units (CFU) per gram of stomach (CFU/gram stomach) for WT and Δ*cagA* after infection in *Mist1-Kras* mice with GIM for 6 weeks. Horizontal line depicts median CFU/g stomach for each strain. **B-D.** CFU/g stomach of WT and Δ*cagA* after co-infection in *Mist1-Kras* mice with GIM for 1 day (**B**), 1 week (**C**), or 6 weeks (**D**). The lines connecting the points distinguish WT (solid symbols) and Δ*cagA* (open symbols) levels from the same mouse. Dotted line represents the limit of detection. **E.** Competitive index scores of B-D. Horizontal line depicts median competitive index score for each timepoint. dpi: days post infection. wpi: weeks post infection. N= 8-17 mice, 2 independent replicates differentiated by symbol shape. Nonparametric two-tailed Mann Whitney U test. ns: nonsignificant.

### CagA promotes Cag-T4SS shutoff after colonization in *Mist1-Kras* mice with GIM, but in a *cagY*-independent manner

GIM stomach tissue has higher levels of baseline inflammation compared to healthy stomach tissue, which could impact Cag-T4SS shutoff (85). Therefore, we tested Cag-T4SS activity for 5-7 output strains from 2-3 mice per experiment (harvested after colonizing the GIM stomach tissue for 6 weeks). As when colonizing wildtype C57BL/6NJ mice, WT output strains had significantly lower levels of IL-8 induction compared to their input controls than Δ*cagA* output strains after colonizing *Mist1-Kras* mice with GIM (median 32% vs. 63%, p-value: 0.0018, nonparametric two-tailed Mann Whitney U test) (**Figure 7A right**). When we compared the levels of IL-8 induction from these output strains to the output strains obtained after 6 weeks of colonization in wildtype C57BL/6NJ mice, WT output strains from *Mist1-Kras* mice with GIM had significantly lower levels of IL-8 induction (median 32% vs. 58%, p-value: 0.0101, nonparametric two-tailed Mann Whitney U test) while the difference between the Δ*cagA* output strains from these timepoints was nonsignificant (median 63% vs. 82%, p-value: 0.18, nonparametric two-tailed Mann Whitney U test) (**Figure 7A**).

**Fig 7.**
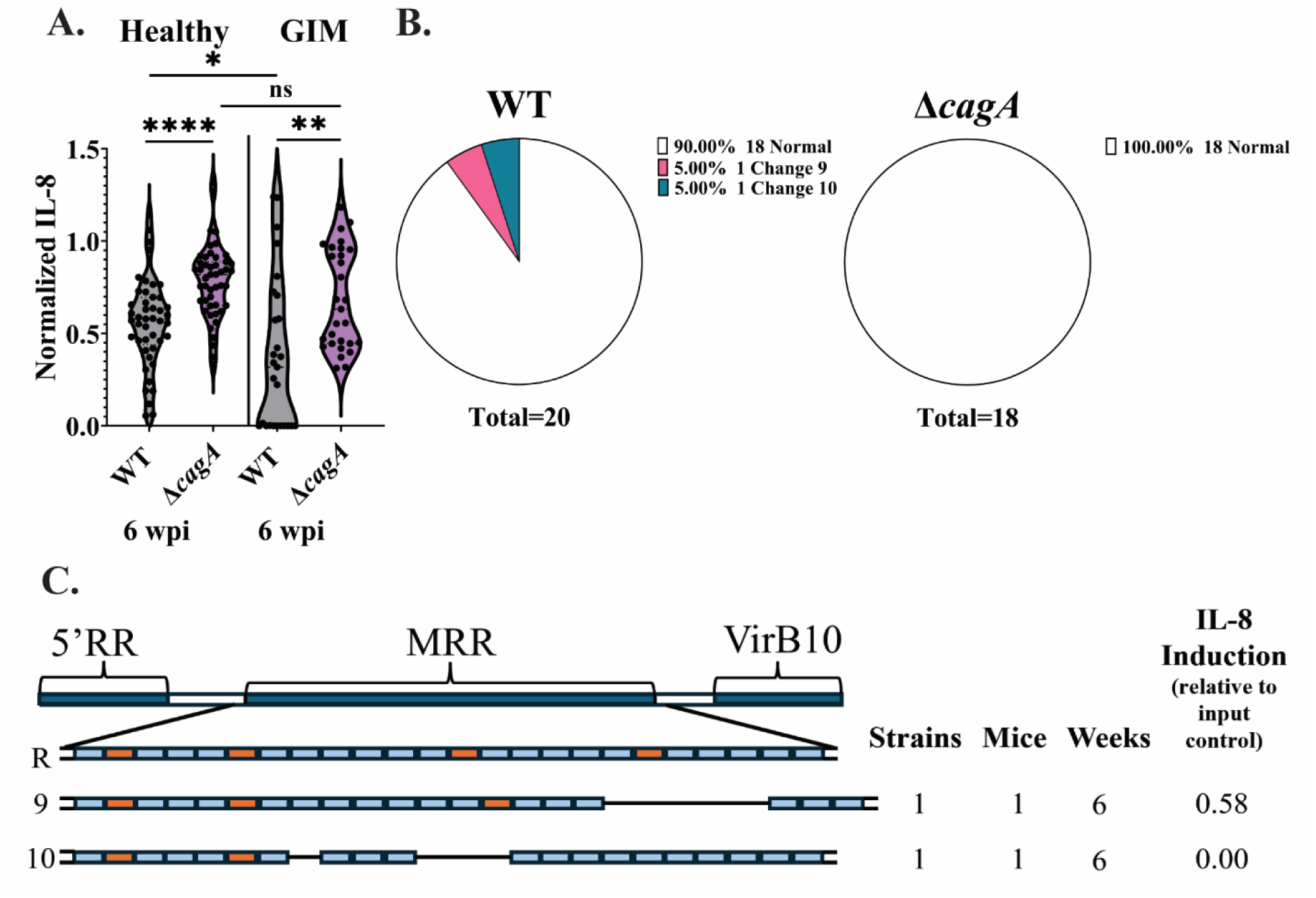
CagA drives Cag-T4SS shutoff during colonization of metaplastic stomach tissue, but *cagY* MRR recombination events are rare. **A.** Levels of IL-8 induction in AGS cells after co-culture with WT or Δ*cagA* output strains after 6 weeks of infection in healthy stomach tissue of wildtype C57BL/6NJ mice (left) or GIM stomach tissue of *Mist1-Kras* mice (right), relative to their respective input controls. Each point represents an individual output strain with 27-47 output strains taken from 4-6 mice from 2 independent experiments per bacterial strain. Nonparametric two-tailed Mann Whitney U test. ns: nonsignificant, *: p-value ≤ 0.05, **: p-value ≤ 0.01, ****: p-value ≤ 0.0001. **B.** Percentage of output strains with normal or changed *cagY* MRR for WT and Δ*cagA* output strains after 6 weeks of infection in *Mist1-Kras* mice with GIM. **C.** Schematic representing the different changes seen to the *cagY* MRR in WT output strains after 6 weeks of infection in *Mist1-Kras* mice with GIM. 5’RR= 5’ repeat region, MRR= middle repeat region, VirB10= *virB10* homology region. Blue= A motifs, Orange= B motifs. R= reference *cagY* MRR. Numbers 9-10 on the left each represent a different change. Strains= the number of different output strains that had the change. Mice= the number of different mice that had output strains with that change. Weeks= weeks postinfection from which output strains were recovered. IL-8 Induction= the relative IL-8 induction output strains had compared to input control.

To see if recombination in the *cagY* MRR was the primary genetic mechanism for attenuating Cag-T4SS activity in strains colonizing the GIM stomach environment, WT and Δ*cagA* output strains with a range of IL-8 induction were chosen for *cagY* MRR sequencing. Despite 11/27 of output strains having complete Cag-T4SS shutoff (having induced less than 10% of the IL-8 seen in the WT input control), only 2/20 WT output strains sequenced had changes in their *cagY* MRR. Again, none of the 18 Δ*cagA* output strains sequenced had changes in *cagY* MRR (**Figure 7B**).

The two WT output strains that had changes in their *cagY* MRR came from the same mouse; however, they had variability in the amount of IL-8 induced from AGS cells during co-culture along with distinct *cagY* MRR changes. The output strain with Change #9 had partial shutoff of the Cag-T4SS (Relative IL-8 induction: 58%) and had an extra A motif along with a separate 438 bp in-frame deletion resulting in the loss of four A motifs and one B motif. The other output strain with Change #10 had completely shut off the Cag-T4SS (Relative IL-8 induction: 0%) and had recombination events resulting in two different in-frame deletions (114 bp and 435 bp, respectively) in the MRR resulting in the loss of three A motifs and one B motif (**Figure 7C**). Similar to our observations in wild-type C57BL/6NJ mice (**Figure 4D**), we did not see any pattern between the *cagY* MRR change size or repeat type altered and the degree of Cag-T4SS shut off.

To determine the *cagY-*independent mechanisms of attenuating Cag-T4SS activity that may be occurring after colonization in *Mist1-Kras* mice with GIM, we again performed WGS on output strains (summarized in **Table 3**). For WT strains, we sequenced one output strain (MsR WT) that did not have a difference in IL-8 induction compared to input (Relative IL-8 induction: 124%) and the only difference detected in the *cag-*PAI was the presence of 3 *cagA* copies. We also sequenced three output strains that had altered IL-8 induction but did not have any differences in the *cagY* MRR. MsS WT (Relative IL-8 induction: 57%) had one copy of *cagA,* a non-synonymous SNP in *cagX* encoding a VirB9 homologue required for Cag-T4SS activity, and a frameshift at the 3’ end of *cagT* (VirB7). MsT WT (Relative IL-8 induction: 26%) had one copy of *cagA* and a different frameshift at the end of *cagT.* Output strain MsU WT (Relative IL-8 induction: 0%) had the same frameshift at the end of *cagT* as MsT WT and a nonsense mutation in *cagI*, encoding CagI which is required for Cag-T4SS activity (18, 88). We also sequenced two output strains from the same mouse where both had lost Cag-T4SS activity (0% IL-8 induction for both strains compared to input control); however, one had a change in the *cagY* MRR while the other did not. MsV WT-1 had Change #10 in the *cagY* MRR and that was the only difference we detected in the *cag-*PAI with WGS. Meanwhile, MsV WT-2 had three copies of *cagA* and a nonsynonymous SNP in the VirB10 homology region of *cagY*.

**Table 3:** Alterations to *cag*-PAI genes found in output strains after six weeks of infection in *Mist1-Kras* mice with GIM.

| Output strain | Locus | Vir gene homology | Polymorphism | Type | IL-8 Induction<sup>a</sup> | CagY MRR<sup>b</sup> |
| --- | --- | --- | --- | --- | --- | --- |
| WT output strains |  |  |  |  |  |  |
| MsR WT | cagA |  | ΔcagA_1 | Copy number variation | 1.24 | NC |
| MsS WT | cagA |  | ΔcagA_1_cagA_2_cagA_3 | Copy number variation | 0.57 | NC |
|  | cag3 |  | R221R (AGA->AGG) | Synonymous SNP |  |  |
|  | cagX | VirB9 | A27E (CGT->CTT) | Non-synonymous SNP |  |  |
|  | cagT | VirB7 | Δ1bp coding (775/843) | Frameshift |  |  |
| MsT WT | cagA |  | ΔcagA_1_cagA_2_cagA_3 | Copy number variation | 0.26 | NC |
|  | cagT | VirB7 | Δ1bp coding (780/843) | Frameshift |  |  |
| MsU WT | cagT | VirB7 | Δ1bp coding (780/843) | Frameshift | 0.00 | NC |
|  | cagI |  | E360* (CTT->ATT) | Nonsense |  |  |
| MsV WT-1 | cagY | VirB10 | MRR variation | Repeat recombination | 0.00 | Change 10 |
| MsV WT-2 | cagA |  | ΔcagA_1 | Copy number variation | -0.03 | NC |
|  | cag3 |  | R221R (AGA->AGG) | Synonymous SNP |  |  |
|  | cagY | VirB10 | A1785D (CGG->CTG) | Non-synonymous SNP |  |  |
| ΔcagA output strains |  |  |  |  |  |  |
| MsW ΔcagA-1 |  |  | Only has ΔcagA_1-4, same as input |  | 1.18 | NC |
| MsX ΔcagA | cagY | VirB10 | +1bp coding (3255/5853) | Frameshift | 0.63 | NC |
|  | cagX | VirB9 | N11N (TTG->TTA) | Synonymous SNP |  |  |
|  | cagW<sup>c</sup> | VirB6 | Δ1bp coding<br>(10/759) | Frameshift |  |  |
|  | cagS |  | +2bp coding<br>(162/600) | Frameshift |  |  |
| MsW<br>Δ*cagA*-<br>2 | cagα | VirB11 | Δ1bp coding<br>(720/993) | Frameshift | 0.42 | NC |
|  | cagα | VirB11 | Δ1bp coding<br>(879/993) | Frameshift |  |  |
|  | cagY | VirB10 | GT→C coding<br>(5503/5853) | Frameshift |  |  |
| MsV<br>Δ*cagA* | cagα | VirB11 | Δ1bp coding<br>(720/993) | Frameshift | 0.40 | NC |
|  | cagU |  | +1bp coding<br>(490/657) | Frameshift |  |  |
| MsU<br>Δ*cagA* | cagT | VirB7 | Δ2bp coding<br>(778/843) | Frameshift | 0.31 | NC |

<sup>a</sup>IL-8 induction relative to respective input controls.
<sup>b</sup>Results of *cagY* MRR sequencing. NC: no change, *cagY* MRR was the same as input control.
Change #: the change seen from *cagY* MRR sequencing (**Figure 7C**).
<sup>c</sup>Annotated as *cagV* in reference sequence (Accession: CP018823); however, sequence homology
is better matched to *cagW* so we have labeled as such.

For the Δ*cagA* output strains, we sequenced one output strain (MsW Δ*cagA*-1) that did not have a difference in IL-8 induction (Relative IL-8 induction: 118%), and we did not see any detectable differences in the *cag-*PAI between this output strain and the input control. We also sequenced four output strains with mid-to-low IL-8 induction (Relative IL-8 induction: 63%-31%) that did not have changes in the *cagY* MRR. MsX Δ*cagA* (Relative IL-8 induction: 63%) had a frameshift mutation in the VirB10 region of *cagY,* a frameshift in *cagW* (VirB6), and a frameshift in *cagS*, a conserved *cag-*PAI gene which encodes the cytosolic protein CagS with unknown function which has not been found to impact Cag-T4SS activity (26). MsW Δ*cagA*-2 (Relative IL-8 induction: 42%) had two different frameshift mutations at the end of *cagα*, encoding a VirB11 homologue required for Cag-T4SS activity, along with a frameshift mutation in the VirB10 region of *cagY.* MsV Δ*cagA* had one of the same frameshifts in *cagα* as MsW Δ*cagA*-2 and a frameshift towards the end of *cagU*, encoding CagU which is required for Cag-T4SS activity (8, 89). Lastly, MsU Δ*cagA* (Relative IL-8 induction: 31%) had a frameshift toward the 3’ end of *cagT.* Overall, we were able to find a variety of alterations in *cag-*PAI genes from WT and Δ*cagA* output strains after infection in *Mist1-Kras* mice with GIM for 6 weeks, with some output strains having changes in multiple *cag-*PAI genes. In this context, we only found a single output strain with *cagY* MRR variation. This suggests that *H. pylori* infecting the GIM stomach environment in *Mist1-Kras* mice may not be manipulating Cag-T4SS activity through *cagY* MRR changes, but rather by altering multiple *cag-*PAI genes, which could have consequences on the Cag-T4SS’s ability to toggle on and off during infection.

## DISCUSSION

CagA and the Cag-T4SS have long been recognized as virulence factors that increase risk for severe disease (ulcer and cancer), but many strains lack the *cag*-PAI, suggesting neither factor is required for colonization or persistent infection. Prior work demonstrated a fitness cost for Cag-T4SS activity after the onset of adaptive immunity that becomes more pronounced with time (56, 65), suggesting that the proinflammatory activities of CagA and the bacterial metabolites translocated by an active Cag-T4SS must be finely tuned to produce the ideal amount of inflammation to promote infection but avoid immune clearance. Consistent with a fitness cost, we observed a small but measurable colonization advantage for a Δ*cagE* Cag-T4SS mutant during single strain infection at six weeks. Furthermore, the intermediate (and not statistically significant) phenotype for the Δ*cagA* mutant indicates that translocation of the CagA effector does not fully explain the Δ*cagE* Cag-T4SS mutant phenotype, suggesting contributions of other substrates. To better detect fitness differences, we used a competitive infection model. Here, we unexpectedly observed a colonization advantage for encoding the Cag-T4SS that manifests as early as one day post-infection with almost complete elimination of Δ*cagE* by six weeks. In the competition setting, the Δ*cagA* mutant displayed complex behavior, showing a subtle advantage at one day, a significant nearly one log deficit at one week, and a return to a colonization advantage in a subset of animals at six weeks. These results support a fitness advantage for *H. pylori* encoding the Cag-T4SS when establishing colonization within the stomach and the importance of other substrates (or activities) besides CagA. This fitness advantage during initial colonization could explain why the *cag*-PAI has been retained in all *H. pylori* lineages except the ancestral HpAfrica2 (67).

Since the initial colonization advantage appeared less dependent on CagA, we hypothesized it resulted from differential proinflammatory cytokine production downstream of host innate immune sensors. However, the colonization advantage persisted even when the known recognition pathways for major proinflammatory bacterial metabolites delivered through the Cag-T4SS were mutated: NOD1 (sensor of peptidoglycan cell wall) and TIFA (sensor of LPS precursors). We also tested a condition with preexisting stomach inflammation (*Mist1-Kras* GIM model) and again found a colonization advantage for strains encoding the Cag-T4SS. In this context, CagA provided no colonization benefit at any timepoint. This model relies on constitutive Ras activity in a subset of gastric epithelial cells, leading to mitogen activated protein kinase (MAPK) signaling, a pathway also activated by translocated CagA. Thus, the selective advantage in strains encoding the Cag-T4SS likely results from the ability to target epithelial cells (rather than immune cells) through CagA-dependent activation of a pathway also activated by Ras signaling, rather than through proinflammatory innate immune receptor pathways.

Both single strain and competitive infection phenotypes became less distinct between six and twelve weeks, which we hypothesize is due to attenuation of Cag-T4SS activity in this timeframe by genetic variation in the *cag-*PAI. Analysis of Cag-T4SS shutoff in our experiments revealed the presence of *cagA* as a strong bacterial driver of shutoff, decreasing Cag-T4SS activity with increased time of infection and presence of GIM (both conditions that increase inflammation).

We used a combination of targeted and whole genome sequencing to gain new insights into the mechanisms of Cag-T4SS attenuation under different conditions. At both six- and twelve-weeks post infection in wildtype C57BL/6NJ mice, we observed frequent alterations in the *cagY* middle repeat region (MRR). Similar to other studies, we could not discern obvious patterns for how these alterations affected activity as both large and small deletions of MRR repeats could result in large or small effects on Cag-T4SS activity (56). Whole genome sequencing confirmed the absence of other mutations in the *cag*-PAI for several *cagY* MRR changes yielding attenuation. Structural modeling suggests that CagY oligomerizes to form a pore spanning the cell surface, outer membrane, and inner membrane (16, 63, 90). Recent structural modeling proposed that sequence variation in CagY influences the Cag-T4SS pore diameter (90, 91) suggesting merit for future structural analyses of these functional variants. Changes to CagY may be attenuating the Cag-T4SS’s ability to translocate CagA while still being able to transport smaller substrates, like the bacterial metabolites, into gastric epithelial cells explaining why recombination in the *cagY* MRR was not found in the output strains lacking *cagA*.

At six weeks post infection in wildtype C57BL6/NJ mice, WT output strains had a recurrent mutation (Change #1) with very mild Cag-T4SS attenuation (50-70% of input) in several mice, raising the possibility that this variant preexisted in the inoculum. This is consistent with observations in the literature that *cagY* variation occurs at a high degree even in *in vitro* culture (56). With increased inflammation at twelve weeks, we observed a higher percentage of MRR mutations, but the variants observed were unique in each animal and one animal even had two distinct mutations. The unique spectrum of mutations raises the possibility of *de novo* recombination events or selective bottlenecks in which only one favorable variant emerges. In the six-week GIM infections, we saw very few *cagY* MRR rearrangements and instead whole genome sequencing revealed mutations, often frameshifts, in other Cag-T4SS structural genes. Many of these completely Cag-T4SS-attenuated outputs had multiple mutations within the *cag*- PAI, which may make it harder to turn Cag-T4SS activity on compared to the strains that only have variation within the MRR of *cagY* or mutations in a single gene in the *cag-*PAI. In WT outputs, we occasionally observed reduction in *cagA* copy number, which also likely contributes to lower IL-8 production. Based on these observations, we propose that recombination-mediated repeat changes may accumulate during liquid culture, but during initial colonization of the stomach, Cag-T4SS activity and CagA translocation promote infection such that preexisting Cag-T4SS attenuated mutants get purged from the population. As infection progresses and inflammation increases, translocation of CagA begins to impart a fitness cost that selects for attenuation of Cag-T4SS activity. Recombination-mediated repeat changes in *cagY* provide a rapid and reversible path to attenuation. Mapping of clones during stomach colonization has revealed that individual bacteria likely seed individual glands that spread and resist superinfection (92). Thus, reactivation of the Cag-T4SS may be required for spreading into new glands. As the stomach becomes further inflamed, strains that have fully attenuated the Cag-T4SS through other mutations increase in frequency even though disfavored during initial colonization.

Together, our results reveal a role for CagA and the Cag-T4SS in promoting stomach colonization, likely through direct interactions with epithelial cells and activation of growth factor-related pathways (MAPK) rather than innate immune pathways (NOD1 and TIFA). At later timepoints the inflamed stomach makes CagA translocation disadvantageous. Further elucidation of the CagA-dependent pathways that both promote and limit infection at different stages of infection could illuminate new host targeted strategies for interception of disease progression and treatment of chronic *H. pylori* infection.

## MATERIALS AND METHODS

### Ethics statement

Authors had access to the study data and were able to review and approve the final manuscript. All mouse experiments were performed in accordance with the recommendations in the National Institutes of Health Guide for the Care and Use of Laboratory Animals. The Fred Hutchinson Cancer Center is fully accredited by the Association for Assessment and Accreditation of Laboratory Animal Care and complies with the United States Department of Agriculture, Public Health Service, Washington State, and local area animal welfare regulations. Experiments were approved by the Fred Hutch Institutional Animal Care and Use Committee, protocol number 1531.

### *H. pylori* strains and culture

*H. pylori* pre-mouse adapted Sydney Strain 1 (PMSS1), referred to as WT in the text, has been previously described (71). Δ*cagE* PMSS1 (Δ*cagE*) has also been previously described (93, 94). All four copies of *cagA* were removed from PMSS1 to generate Δ*cagA* PMSS1 (Δ*cagA*) as previously described (78). WT and derivatives (Δ*cagE* and Δ*cagA*) were grown on agar containing 4% Columbia agar (Oxoid), 5% defibrinated horse blood (Hemostat Labs), 0.2% β-cyclodextrin (Thermo Fisher), 10 μg/mL vancomycin (Thermo Fisher), 2.5 U/mL polymyxin B (Sigma-Aldrich), and 8 μg/mL amphotericin B (Sigma-Alrdrich). Strains were grown in liquid media containing 90% (vol/vol) Brucella broth (BD BBL, Fisher) and 10% heat-inactivated fetal bovine serum (FBS) (Innovative Bioscience), referred to as BB10. For mouse infections, bacteria were grown on agar plates, used to inoculate fresh BB10 and were cultured shaking overnight. Cultures were grown to an optical density at 600 nm (OD600) of 0.4-0.6 (mid-log phase), from which an inoculum of ∼5 x 10^7^ CFU/100 μL BB10 was prepared. To determine *H. pylori* titers in the mouse stomach, harvested tissue was weighed, homogenized, serially diluted, and plated on the agar media as described above with the addition of 5 μg/mL cefsulodin (Thermo Fisher), 5 μg/mL trimethoprim (Sigma), and 200 μg/mL bacitracin (Aros Organics, Fisher) to prevent growth of contaminating stomach microbiota. For competition experiments, 15 μg/mL of chloramphenicol was added to agar plates to distinguish between cassette-marked (chloramphenicol resistant) Δ*cagE* / Δ*cagA* and unmarked (chloramphenicol sensitive) WT strains. All strains cultured on agar plates and in liquid media grew at 37°C under microaerophilic conditions of 10% CO_2_, 10% O_2_, and 80% N_2_ maintained in a tri-gas incubator.

### Mouse lines and husbandry

All mice were housed in sterilized microisolator cages with irradiated rodent chow, autoclaved corn cob bedding, and acidified, reverse-osmosis purified water.

Wildtype C57BL/6NJ mice were purchased from Jackson Laboratories (strain #005304).

*Mist1-CreERT2^Tg/+^; LSL-K-Ras(2D)^Tg/+^* mice (*Mist1-Kras*) were bred as previously described (85). Ear punches were collected and used for genotyping as previously described (85, 86).

*Tifa* knock out mice (*Tifa -/-*) (95) were from Dr. Scott D. Gray-Owen. Ear punches were collected and used for genotyping with the following primers: *Tifa* F: CTTCCCTCTGCTTCCCCTAC, *Tifa* R: TTCTTTGTAGAGCCGAAACTCA. *Nod1* knock out mice (*Nod1-/-*) (96) were originally from Millennium Pharmaceuticals. Ear punches were collected and used for genotyping with the following primers: *Nod1* WT F: CTTAGGCATGACTCCCTCCTGTCG, *Nod1* WT R: GATCTTCAGCAGTTTAATGTGGGAGTGAC, *Nod1* KO F: CTTAGGCATGACTCCCTCCTGTCG, *Nod1* KO R: CCATTCAGGCTGCGCAACTGTTG.

*Tifa/Nod1* mice were bred by crossing *Tifa -/-* and *Nod1 -/-* mice. Ear punches were collected and used for genotyping with the following primers: *Tifa* F: CTTCCCTCTGCTTCCCCTAC, *Tifa* R: TTCTTTGTAGAGCCGAAACTCA, *Nod1* WT F: CTTAGGCATGACTCCCTCCTGTCG, *Nod1* WT R: GATCTTCAGCAGTTTAATGTGGGAGTGAC, *Nod1* KO F: CTTAGGCATGACTCCCTCCTGTCG, *Nod1* KO R: CCATTCAGGCTGCGCAACTGTTG.

### Animal Infection Experiments

For colonization experiments in healthy stomach tissue, 8–10-week-old wildtype C57BL/6NJ female and male mice were orally administered a mid-log *H. pylori* culture of ∼5x10^7^ CFU in 100 μL of BB10, via oral gavage. For competition experiments, mice were infected with a 50:50 mixture of WT and Δ*cagE* or Δ*cagA*, totaling ∼5x10^7^ CFU in 100 μL of BB10.

For colonization experiments using *Tifa* and/or *Tifa/Nod1* mice, 6–83-week-old female and male mice were orally administered a mid-log *H. pylori* culture of ∼5x10^7^ CFU in 100 μL of BB10, via oral gavage. For competition experiments, mice were infected with a 50:50 mixture of WT and Δ*cagE* or Δ*cagA*, totaling ∼5x10^7^ CFU in 100 μL of BB10.

For colonization experiments in *Mist1-Kras* mice, Kras^G12D^ expression in *Mist1+* cells was induced in 7–17-week-old female and male mice by three subcutaneous doses of 5 mg of tamoxifen in corn oil over 3 consecutive days. Metaplastic progression via Kras^G12D^ expression was allowed to develop for 6 weeks, a timepoint which we refer to as gastric intestinal metaplasia (GIM). At this timepoint, mice were orally administered a mid-log *H. pylori* culture of ∼5x10^7^ CFU in 100 μL of BB10, via oral gavage. For competition experiments, mice were infected with a 50:50 mixture of WT and Δ*cagE* or Δ*cagA*, totaling ∼5x10^7^ CFU in 100 μL of BB10.

Mice were humanely euthanized via CO_2_ inhalation followed by cervical dislocation. Stomachs were aseptically harvested, cutting most of the forestomach away to leave only glandular epithelium. Stomachs were then cut along the degree of lesser curvature and stomach contents were removed by lightly scraping the luminal side. Stomachs were then cut in half, with one half being weighed, homogenized, and serially diluted to determine titers as described earlier.

### AGS co-culture and IL-8 ELISA assays

Output strains were recovered from infected mice after euthanasia through serial dilution plating for CFU, described above. Up to 12 individual colonies per strain per mouse were expanded and frozen at -70℃ in freeze media containing Brain Heart Infusion Media+ (BD Bacto), 10% FBS (Innovative Bioscience), 20% Glycerol (Thermo Scientific) and 0.2% cyclodextrin (Thermo Fisher). These output strains were then grown on agar plates and used for liquid cultures (described above).

Concurrently, approximately 100,000 human AGS gastric adenocarcinoma cells (ATCC: CRL-1739) per well were seeded in 24-well plates in DMEM (Gibco) with 10% FBS (Innovative Bioscience) (DMEM10) and allowed to grow overnight at 37°C with 10% CO_2_ and 10% O_2_. The next day, media was removed and replaced with infection media (containing 80% DMEM10 and 20% BB10) inoculated with *H. pylori* output strains for an MOI of 10:1. Infection media with no *H. pylori* added was used as a control. The input strain for WT and Δ*cagA* were used as positive and baseline input controls. Δ*cagE* was used as a negative control. These controls were included for each set of AGS co-culture experiments. Infections were performed in duplicate wells. Supernatants were harvested 24 hours after co-culture and stored at -20°C. Prior to the IL-8 ELISA assay, supernatants were thawed and diluted 1:4 in assay diluent (BioLegend).

IL-8 in the supernatant was quantified using the ELISA MAX Deluxe Set Human IL-8 kit (BioLegend) according to the manufacturer’s protocol. Samples were run in triplicate. To account for variability in the assay, IL-8 values were normalized to the respective input strain present on each ELISA plate and measured concurrently. **Supplemental Figure S4** contains examples of the raw IL-8 (pg/mL) produced by the AGS cells after co-culture with the output strains and controls.

### *cagY* PCR and sequencing

Reactions were performed in a total volume of 50 μL containing 10 ng of genomic DNA, 200 μM of each dNTP, 0.5 μM of each primer (F: *5’-CCGTTCATGTTCCATACATCTTTG-3’*, R: *5’-CTATGGTGAATTGGAGCGTGTG-3’*), 0.02 U/μL Phusion DNA polymerase, and 1x Phusion Buffer (Thermo Scientific). PCR products were purified using the Monarch Spin PCR & DNA Cleanup Kit (New England Biolabs), diluted to be within 20-200 ng/μL, and sent for standard purified amplicon sequencing through Plasmidsaurus.

### Oxford Nanopore Whole Genome Sequencing and Analysis

DNA extraction, purification, and sequencing were done as previously described (97). High quality *H. pylori* genomic DNA was extracted with a CsCl gradient to prevent shearing. Two full plates of healthy mid-log bacteria were scraped and resuspended in 3.5 mL of STE (0.1 M NaCl, 10 mM Tris-HCl, 1 mM EDTA, pH 8.0) with 350 μg/mL of lysozyme (Fisher), then incubated at 37 ⁰C for 15 minutes. Complete lysis was performed by adding 350 µL of 10% SDS (Fisher Bioreagents) and incubating at 65 ⁰C for 15 minutes. Proteins were digested with 35 µL of proteinase K (50 mg/mL) (Invitrogen) at 50 ⁰C for 2.5 hours. Solid CsCl (Sigma-Aldrich) was added to a final concentration of 1 g/mL, mixed by inversion, then incubated at 65 ⁰C for 15 minutes. Ethidium bromide (10 mg/mL) (Fisher Biotech) was added at 80 µg/mL, then incubated at 65 ⁰C for 40 minutes. Samples were transferred to g-Max Quick-Seal tubes (Beckman Coulter), sealed, and centrifuged at 390,000 g (Beckman Coulter - Type 70.1Ti rotor) at 20 ⁰C overnight. DNA was visualized with UV and carefully extracted from the tubes with a 20-gauge needle and syringe. DNA was then cleaned via butanol extraction at least five times with one volume of STE-saturated butanol. DNA was then precipitated with five volumes of 77% ethanol. DNA strands were transferred to 200 μL of TE (10 mM Tris-HCl, 100 mM EDTA, pH 8.0) for storage using flame-closed glass Pasteur pipettes and allowed to rehydrate at 4 ⁰C overnight before freezing. Samples were quantified and quality checked by Tapestation (Agilent 4200), then submitted for standard bacterial genome sequencing through Plasmidsaurus. Genome assemblies from Plasmidsaurus were aligned to the PMSS1 reference sequence (Accession: CP018823) using Mauve alignment in Geneious. Polymorphisms in *cag-*PAI genes were noted.

## Data Availability

Sequencing data produced in this study has been deposited in the NCBI (BioProject: PRJNA1507379).

## Acknowledgements

This work was supported by NIH R01 AI054423 (NRS), NIH/NCI R00 CA263036 (VPO), the Canada Research Chairs Program (SDG), Canadian Institutes of Health Research Project (grant #197806) (SDG), and from the NIH P30 CA015704 of the Fred Hutch/University of Washington/Seattle Children’s Cancer Consortium, which includes the Comparative Medicine Shared Resource, RRID: SCR_022610.

## Author Contributions

Based on Contributor Role Taxonomy (CRediT).

**JAS:** Conceptualization, Data Curation, Formal Analysis, Investigation, Methodology, Project Administration, Validation, Visualization, Writing - Original Draft, Writing - Review & Editing; **JPF:** Data Curation, Formal Analysis, Investigation, Methodology, Writing - Review & Editing; **VPO:** Funding Acquisition, Investigation, Methodology, Writing-Review & Editing; **CXG:** Resources, Writing-Review & Editing; **SDG:** Funding Acquisition, Resources, Supervision, Writing-Review & Editing; **NRS:** Conceptualization, Data Curation, Funding Acquisition, Methodology, Project Administration, Resources, Supervision, Writing –Original Draft, Writing-Review & Editing.

